# Small molecule resensitizes *Salmonella* to itaconate and decreases bacterial proliferation in macrophages

**DOI:** 10.1101/2020.07.15.204305

**Authors:** Dustin Duncan, Justin H. Chang, Maxim N. Artyomov, Karine Auclair

**Affiliations:** Department of Chemistry, McGill University, Montreal, Quebec, Canada; Department of Pathology and Immunology, Washington University in St. Louis, St. Louis, Missouri, USA

**Keywords:** glyoxylate, intracellular, isocitrate lyase, itaconate, macrophage, resensitize, *Salmonella*

## Abstract

Antimicrobial resistance is a global health crisis offering little reprieve. The situation urgently calls for new drug targets and therapies for infections. We have previously suggested a different approach to treat infections, termed bacterio-modulation, in which a compound modulates the bacterial response to the host immune defense. Herein we show that monocytes infected with *Salmonella enterica* spp. Typhimurium can be cured using non-antimicrobial compounds that resensitize the bacterium to itaconate, a macrophage-derived antimicrobial metabolite. We propose that this represents a novel strategy to treat infections.

## Introduction

Antimicrobial resistance continues to spread, rapidly putting out of use pharmaceuticals that took decades to develop. The perpetual arms race between humans and bacteria warrants further exploration of bacterial targets for the development of new antimicrobial therapies. Among the pathways that have been investigated is the glyoxylate cycle.^1-2^ This pathway is particularly attractive because it is *conditionally* necessary for survival and absent in mammals, thus limiting resistance selection and off-target effects. Via the two enzymes isocitrate lyase (Icl) and malate synthase (Ms), the glyoxylate cycle is indispensable to bacteria in carbon-starved environments and avoids the loss of carbon atoms through decarboxylation (Figure 1). This is supported by experiments in which the pathway is rendered dysfunctional in pathogenic bacteria, resulting in a loss or a significant reduction in pathogenicity.^3-6^ Targeting this pathway is expected to only affect the pathogenic bacteria in nutrient-starved milieus, such as within the macrophage, while the commensal bacteria, often found in nutrient-rich locations, should remain unaffected, thus treating infections with minimal effect on the beneficial microflora. Whereas several research groups have attempted to target Icl for the development of antibiotics,^7-14^ such inhibitors have yet to make it to the market.

**Figure 1:**
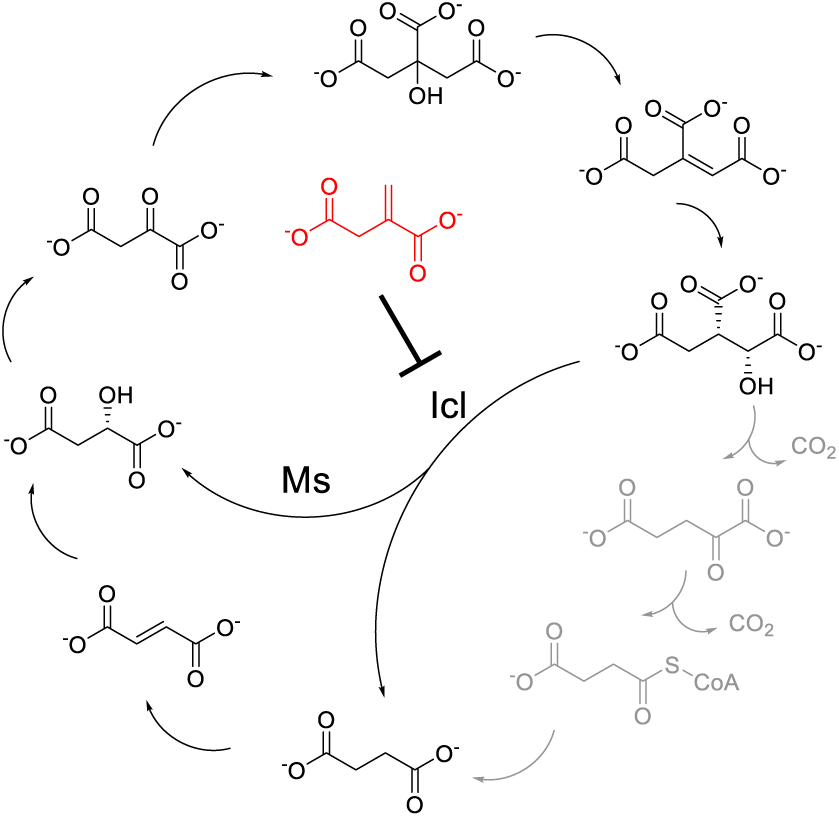
The glyoxylate cycle (black) as it relates to the Kreb’s cycle. Itaconate (red), produced by Irg1 from *cis*-aconitate, inhibits Icl, the first of two enzymes that divert D-isocitrate to succinate and L-malate.

Nature has evolved to take advantage of the glyoxylate cycle as an antimicrobial target. Upon sensing lipopolysaccharides from bacteria, macrophages begin reprogramming their metabolism and produce the antimicrobial metabolite itaconate, a known inhibitor of Icl (Figure 1).^15-16^ The immunoresponsive gene 1 (*irg1*), which encodes for the itaconate-producing protein Irg1, or *cis*-aconitate decarboxylase (CAD or Acod1), is among the most upregulated genes during activation of pro-inflammatory macrophages and microglial cells.^17-18^ Irg1 catalyses the decarboxylation of the Kreb’s cycle intermediate *cis*-aconitate to produce itaconate. Despite the production of the antimicrobial itaconate by activated macrophages, some intracellular pathogens such as *Mycobacterium tuberculosis, Salmonella enterica, Yersinia pestis*, and *Pseudomonas aeruginosa* can thrive in macrophages in part due to a pathway dedicated to transforming itaconate to products that feed back into the bacterial Kreb’s cycle.^19^

We report herein that compound **1** which is known to resensitize *Salmonella enterica* spp. Typhimurium to itaconate,^20^ is also successful at preventing bacterial proliferation in macrophages via an itaconate-dependent mechanism.

## Materials and Methods

### General Cell Growth Procedure

BV2 cells were maintained in cell culture dishes (Eppendorf) in Dulbecco’s Modified Eagle’s Medium (DMEM, Gibco) supplemented with 10% heat-inactivated fetal bovine serum (heat denatures pantetheinases) (Gibco), 2 mM L-glutamine (100x Glutamax, Gibco), and 100 U/mL penicillin-streptomycin (Gibco), incubated at 37°C in 5% CO_2_. Cells were divided every 3 days. To propagate the BV2 cells, the growth medium was removed by aspiration and the cells were washed with 3 mL PBS (Gibco), before adding 3 mL of Cell Stripper (Corning). The BV2 cells were incubated with the Cell Stripper for ca. 5-10 minutes at 37°C in 5% CO_2_. After removing the medium, the cells were washed with ca. 5 mL DMEM, diluted into fresh DMEM (10-fold) and re-plated. For the *irg1*^*+/+*^ and *irg1*^*-/-*^ BMDM cells, they were isolated as previously described^21^ from *irg1*^*+/+*^ or *irg1*^*-/-*^ C57BI/6N mice and sustained in RPMI-1640 (Gibco) supplemented with 10% heat-inactivated fetal bovine serum, 2 mM L-glutamine, and 100 U/mL penicillin-streptomycin, and incubated at 37°C in 5% CO_2_.

### Alamar Blue Assay

The 5× resazurin solution was prepared by dissolving resazurin sodium salt (11.4 mg) in PBS (100 mL) and filter sterilized. BV2 or BMDM cells were plated using 100 µL of respective growth medium at a concentration of 2 × 10^5^ cell/mL. The cells were allowed to adhere to the 96-well plates (black, flat-bottom, transparent bottom) for 2 hours, before washing with growth media (100 µL), and incubation with respective growth medium containing **1** or **2** (synthesized as described by Hammerer *et al*.^20^) at concentrations ranging from 0 – 0.5 g/L. The BV2 or BMDM cells were then incubated for 24 hours at 37°C and 5% CO_2_, before addition of 25 µL of 5× resazurin solution and incubation at 37°C in 5% CO_2_ for 30 minutes. Fluorescence measurements were performed using a plate-reader with excitation at 540 nm, and emission at 590 nm.

### *In Cellulo* Assay

This procedure is adapted from Vladoianou *et al*.^22^ BV2 cells (2 × 10^5^ cells/mL, 100 µL) in antibiotic-free DMEM containing 5 mM itaconic acid were added to 96-well plates (black, flat-bottom, transparent bottom) and allowed to adhere for 24 hours at 37°C with 5% CO_2_. From an LB plate with *S*. Typhimurium colonies, 4 colonies were picked and inoculated in 10 mL of LB broth, before incubation at 37°C with shaking at 200 rpm, until an OD_600_ of approximately 0.6 (equal to approximately 5 × 10^8^ CFU/mL) was reached. The *S*. Typhimurium culture was diluted into antibiotic-free DMEM to a concentration of 6 × 10^5^ CFU/mL (i.e. MOI of 30) and incubated with the BV2 cells for 10 minutes at 37°C with 5% CO_2_. The growth medium was removed by aspiration and the BV2 cells were washed with 2 × 100 µL gentamicin-containing (0.200 µg/mL) DMEM. The resulting *S*. Typhimurium-infected BV2 cells were exposed to a mixture of 0.200 µg/mL gentamicin, 5 mM itaconic acid, and **1** or **2** at 0.25, 0.5, or 1.0 g/L, and incubated for 24 hours at 37°C in 5% CO_2_. The growth medium was then removed by decantation and the BV2 cells were washed with 100 µL PBS. The BV2 cells were next lysed by adding 100 µL of 0.1% Triton X-100 at 4°C for 15 minutes. The BV2 cell lysate was plated at 10× and 100× dilutions in PBS onto LB-agar medium and incubated at 37°C for 10 – 11 hours.

## Results

### Toxicity of compounds 1 and 2

Neither **1** nor **2** were found to show antibacterial activity towards *S*. Typhimurium.^20^ We measured the toxicity of both compounds towards BV2 cells at concentrations up to 0.5 g/L (approximately 1.4 mM; higher than the concentration needed to resensitize bacteria to itaconate *in vitro*).^20^ Neither of the compounds affected BV2 cell growth (Figure 2) as determined from the Alamar Blue assay. When testing the toxicity of **1** and **2** on *irg1*^+/+^ BMDM cells, there was an increase in fluorescence in the Alamar Blue assay in a dose-dependent manner for **1**, whereas **2** did not elicit any change in growth (Figure 3). Compound **1** was evaluated for toxicity in *irg1*^-/-^ BMDM cells, and there was no effect on the cells (Figure 4).

**Figure 2.**
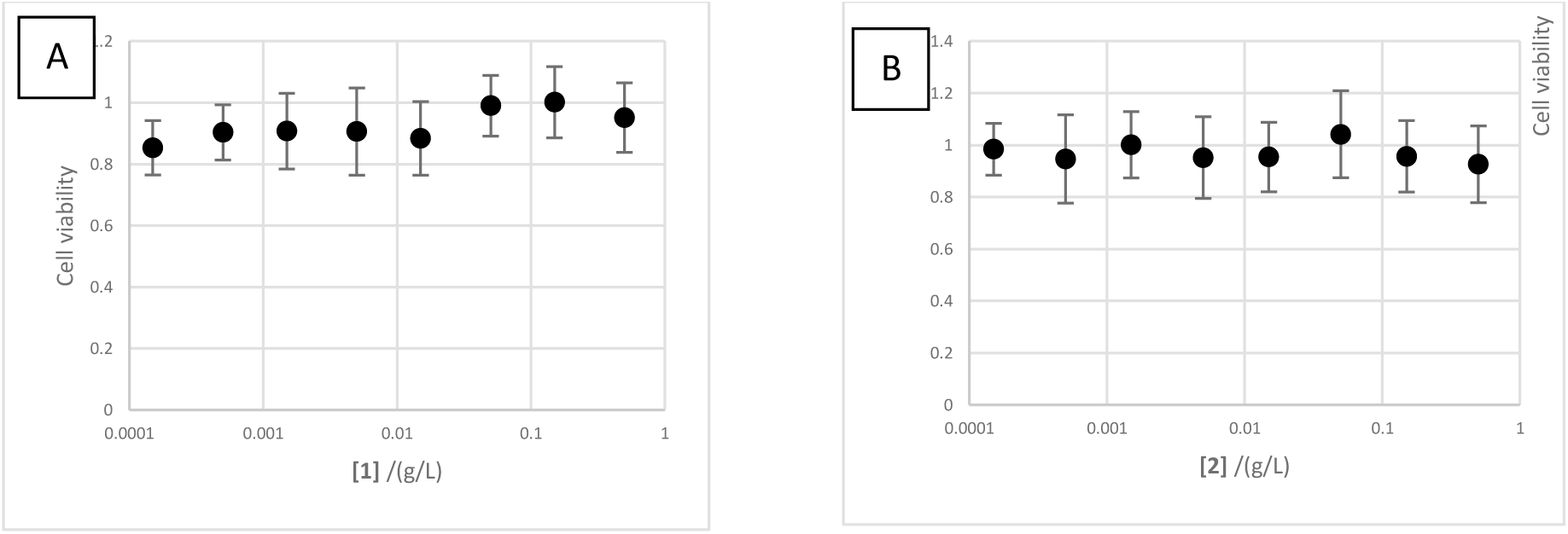
Viability of BV2 cells in the presence of **1** (panel A) or **2** (panel B), as determined using the Alamar Blue assay. Error bars = SEM, n = 3.

**Figure 3:**
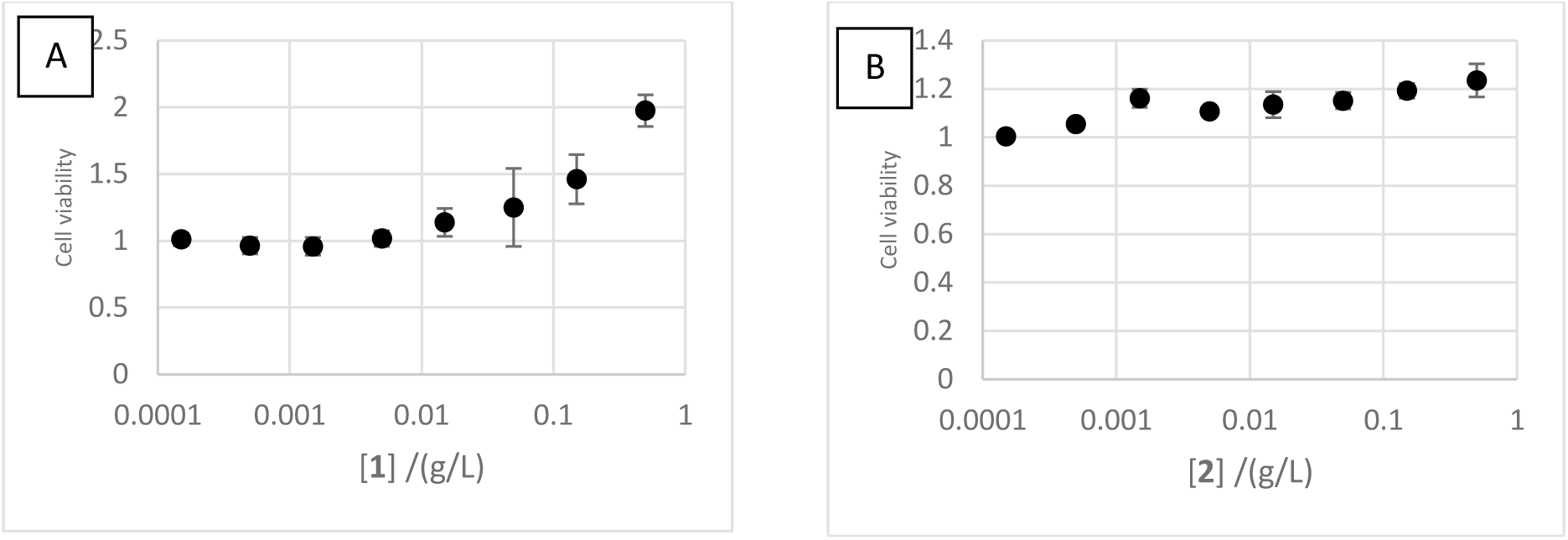
Viability of BMDM *irg1*^*+/+*^ cells as determined using the Alamar Blue assay. Error bars = SEM, n = 3 for **1** (panel A), n = 2 for **2** (panel B)

**Figure 4:**
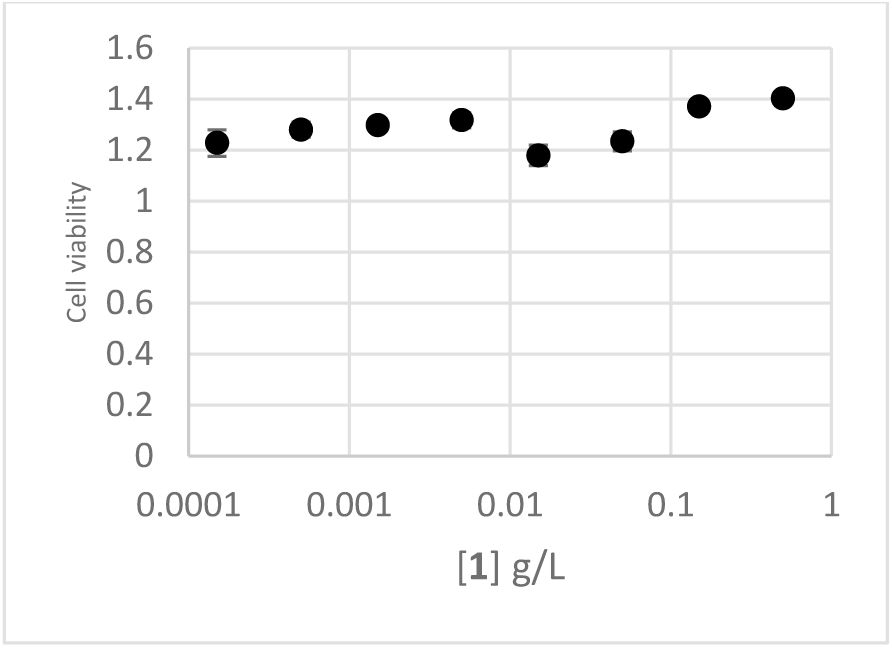
Viability of BMDM *irg1*^*-/-*^ cells in the presence of compound **1** as determined using the Alamar Blue assay. Error bars = Standard deviation for one triplicate experiment

### Efficacy of 1 and 2 at reducing bacterial proliferation in BV2 cells

We evaluated the efficacy of **1** and **2** at reducing the proliferation of *S*. Typhimurium in BV2 cells. The monocytes were incubated with 5 mM itaconic acid overnight for consistency in the abundance of itaconate within the experimental conditions. This concentration is known to recapitulate the effects of endogenous itaconate expression.^23^ After a 10-minute challenge of BV2 cells to *S*. Typhimurium (at a multiplicity of infection or MOI of 30, *i*.*e*. 30 *Salmonella* cells per BV2 cell), compound **1**, compound **2**, or the vehicle alone, was added to the growth medium and the cells were allowed to proliferate for an additional 24 hours. Gentamicin was present in the media to kill extracellular bacterial cells after challenging the BV2 cells with *S*. Typhimurium. The results (Figure 5) demonstrate that in the presence of compound **1**, bacterial proliferation in BV2 cells is reduced by approximately 99.9%, whereas addition of compound **2** or the vehicle did not clear the infection significantly.

**Figure 5.**
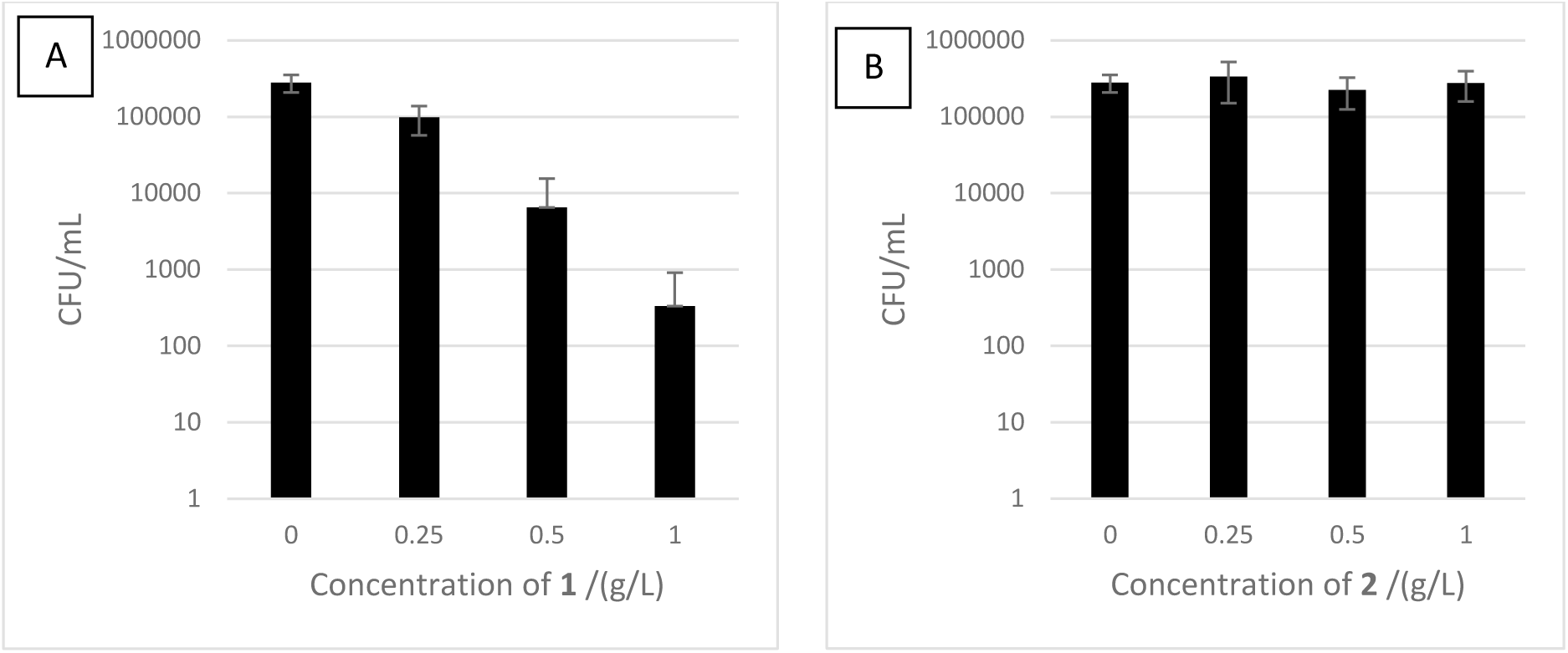
CFU/mL of *S*. Typhimurium isolated from infected BV2 cells incubated with **1** (panel A) or **2** (panel B). Error bars = SEM, n = 3.

## Discussion

BV2 cells were selected for our experiments not only for their phagocytic capacities,^24^ but also because they have been used extensively to study the role of itaconate during infections.^25-26^ The compounds used herein were designed as prodrug inhibitors of itaconate degradation.^20,27^ They are pantothenamide derivatives, selectively transformed into the corresponding coenzyme A (CoA) analogues by three enzymes of the CoA biosynthetic pathway (Figure 6): pantothenate kinase (PanK), phosphopantetheine adenylyltransferase (PPAT), and dephospho-CoA kinase (DPCK). Importantly, although many pantothenamides show antimicrobial activity,^28^ compounds **1** and **2** do not. Once bioactivated, compound **1** resensitizes *S*. Typhimurium to itaconate by inhibiting itaconate-degradation within the bacterium.^19-20^ We opted here to also work with prodrug **2**, which does not resensitize *S*. Typhimurium to itaconate, and serves as a negative control.

**Figure 6.**
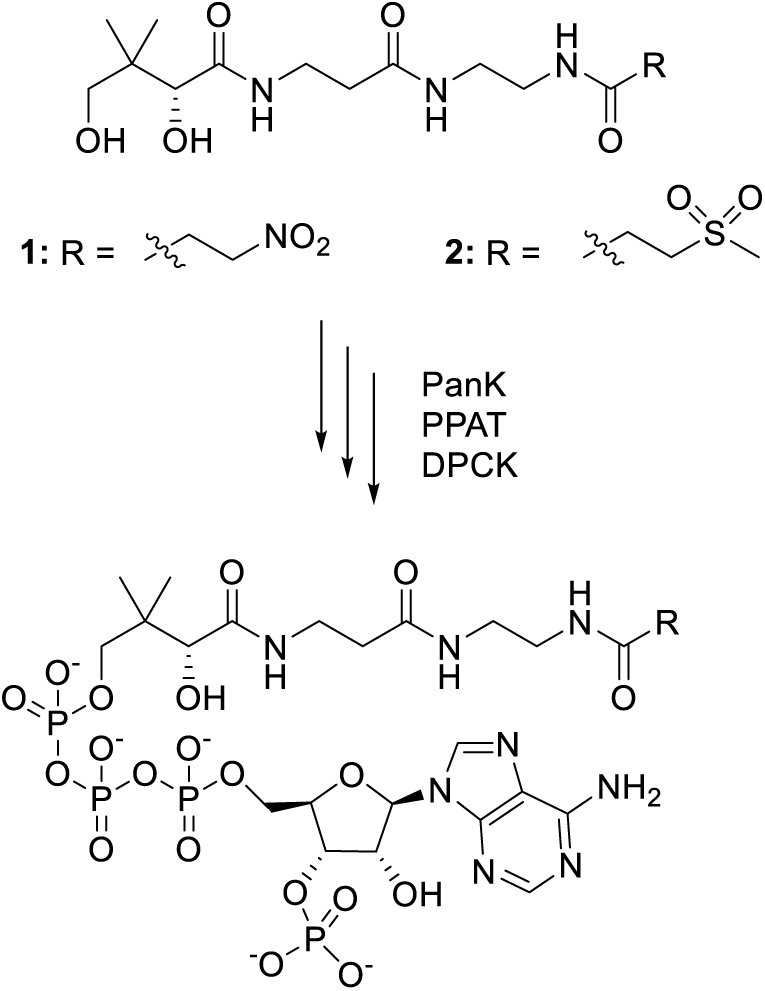
Structure of compounds **1** and **2**, and their bioactivation by the CoA biosynthetic pathways

As seen in Figure 2, there was no change in fluorescence in the Alamar Blue assay for compounds **1** or **2**, and this was interpreted to be no loss of cell viability of BV2 cells, and therefore neither compounds **1** nor **2** are cytotoxic. After infection of the BV2 cells with *S*. Typhimurium in the presence of itaconate and **1**, it was found that the bacterial proliferation was reduced by about 99.9%, whereas treatment with **2** did not elicit a reduction in bacterial proliferation (Figure 5). Remarkably, this is also consistent with compound **1** penetrating both the macrophage and this Gram-negative bacterial strain. These results are in agreement with the *in vitro* experiments showing that **1** can resensitize S. Typhimurium under nutrient-depleted conditions that partially mimic the environment within the phagolysosome, whereas **2** cannot.^20^ Overall these data suggest that it may be possible to treat bacterial infections with such bacterio-modulators instead of antibacterials.

In an attempt to confirm this result with a different model, the same protocol was repeated with bone-marrow derived macrophages (BMDM), encoding Irg1 or not. The toxicity of **1** and **2** on *irg1*^*+/+*^ BMDM cells was examined, and neither compound was found to negatively affect growth (Figure 3). Interestingly **1** lead to a marked dose-dependent increase in signal from the Alamar Blue assay, whereas **2** did not. Since the Alamar Blue assay measures metabolic turnover, this suggests that **1** has some role in increasing aerobic respiration in BMDM cells.

We next performed a preliminary experiment using irg1^-/-^ BMDMs cells (Figure 4) to investigate the role of the *irg1* gene on the BMDM cells’ response to **1**. The effect that **1** had on the *irg1*^*+/+*^ BMDM cells was greatly reduced once *irg1* was knocked out. It is unclear why this is the case considering that *irg1* should not be expressed appreciably given that there was no LPS present to activate its expression. This may provide evidence that **1** may interact with regulatory elements of *irg1* expression that affects respiration. It was not possible to determine the efficacy of **1** or **2** in clearing *S*. Typhimurium infections from BMDM cells due to their intrinsic pyroptosis response, a mechanism of programmed cell death which *in vivo* induces an inflammatory response to recruit other cells.^29^

In summary, the premise of our approach is that non-antibiotic compounds, such as **1**, may not provide significant evolutionary pressures to select for resistant strains because they are neither bacteriostatic nor bactericidal, yet may resensitize bacteria to the natural killing capacity of macrophages, thereby helping to clear infections. Macrophages are superbly tooled to kill bacteria. By producing selective antimicrobials, such as itaconate, that are lethal only under specific conditions, macrophages leave little opportunity for resistance selection. Some (relatively rare) intracellular pathogens have however evolved several mechanisms allowing them to proliferate in macrophages, including the ability to metabolise itaconate to pyruvate and acetyl CoA. Given the efficacy of persisters in undermining typical antibiotic treatments, we propose that resensitizing such pathogens to itaconate may be an especially effective strategy to treat chronic infections.^30^ *S*. Typhimurium proliferates rapidly in BV2 cells. Consistent with the proposed antimicrobial role of itaconate, the addition of compound **1**, designed to inhibit itaconate degradation, reduces bacterial proliferation in BV2 cells by up to 99.9%. Compound **1** is however rapidly hydrolyzed by pantetheinases (vanins) in fresh blood, precluding *in vivo* investigations. Interestingly, several studies have shown that such pantothenamide derivatives can be rendered more stable, while maintaining their biological activity, by chemical modifications at the labile amide,^31-35^ the geminal dimethyl,^36-37^ or the β-alanine moiety,^38-39^ warranting further investigations.

## Acknowledgements

*This work was supported by research grants from the Canadian Institute of Health Research (CIHR) and the Natural Sciences and Engineering Research Council (NSERC) to K*.*A*.

## References

1. Dolan SK. Welch M. The Glyoxylate Shunt, 60 Years On. Annu Rev Microbiol. 2018;72:309–30.

2. Lorenz MC. Fink GR. Life and death in a macrophage: role of the glyoxylate cycle in virulence. Eukaryot Cell. 2002;1:657–62.

3. Munoz-Elias E J. McKinney JD. Mycobacterium tuberculosis isocitrate lyases 1 and 2 are jointly required for in vivo growth and virulence. Nat Med. 2005;11:638–44.

4. McKinney JD. et al. Persistence of Mycobacterium tuberculosis in macrophages and mice requires the glyoxylate shunt enzyme isocitrate lyase. Nature. 2000;406:735–8.

5. Lindsey TL. Hagins JM. Sokol PA. Silo-Suh LA. Virulence determinants from a cystic fibrosis isolate of Pseudomonas aeruginosa include isocitrate lyase. Microbiology. 2008;154:1616–27.

6. Fang FC. Libby SJ. Castor ME. Fung AM. Isocitrate lyase (AceA) is required for Salmonella persistence but not for acute lethal infection in mice. Infect Immun. 2005;73:2547–9.

7. Tiwari A. Kumar A. Srivastava G. Sharma A. Screening of Anti-mycobacterial Phytochemical Compounds for Potential Inhibitors against Mycobacterium Tuberculosis Isocitrate Lyase. Curr Top Med Chem. 2019;19:600–8.

8. Krátký M. et al. Phenolic N-monosubstituted carbamates: Antitubercular and toxicity evaluation of multi-targeting compounds. Eur J Med Chem. 2019;181:111578.

9. Lee YV. Choi SB. Wahab HA. Lim TS. Choong YS. Applications of Ensemble Docking in Potential Inhibitor Screening for Mycobacterium tuberculosis Isocitrate Lyase Using a Local Plant Database. J Chem Inf Model. 2019;59:2487–95.

10. Krátký M. et al. Salicylanilide diethyl phosphates as potential inhibitors of some mycobacterial enzymes. Sci World J. 2014;2014:703053.

11. Yang HC. et al. Synthesis and evaluation of hydroquinone derivatives as inhibitors of isocitrate lyase. Arch Pharm Res. 2007;30:955–61.

12. Ji L. Long Q. Yang D. Xie J. Identification of mannich base as a novel inhibitor of Mycobacterium tuberculosis isocitrate by high-throughput screening. Int J Biol Sci. 2011;7:376–82.

13. Fahnoe KC. et al. Non-traditional antibacterial screening approaches for the identification of novel inhibitors of the glyoxylate shunt in gram-negative pathogens. PLoS One. 2012;7:e51732.

14. McVey AC. et al. 2-Aminopyridine Analogs Inhibit Both Enzymes of the Glyoxylate Shunt in Pseudomonas aeruginosa. Int J Mol Sci. 2020;21:2490.

15. McFadden BA. Purohit S. Itaconate, an isocitrate lyase-directed inhibitor in Pseudomonas indigofera. J Bacteriol. 1977;131:136–44.

16. Honer Zu Bentrup K. Miczak A. Swenson DL. Russell DG. Characterization of activity and expression of isocitrate lyase in Mycobacterium avium and Mycobacterium tuberculosis. J Bacteriol. 1999;181:7161–7.

17. Degrandi D. Hoffmann R. Beuter-Gunia C. Pfeffer K., The proinflammatory cytokine-induced IRG1 protein associates with mitochondria. J Interferon Cytokine Res. 2009;29:55–67.

18. Thomas D.M. Francescutti-Verbeem D.M. Kuhn D.M. Gene expression profile of activated microglia under conditions associated with dopamine neuronal damage. FASEB J. 2006;20:515–7.

19. Sasikaran J. Ziemski M. Zadora P.K. Fleig A. Berg I.A. Bacterial itaconate degradation promotes pathogenicity. Nat Chem Biol. 2014;10:371–7.

20. Hammerer F. Chang J.H. Duncan D. Castaneda Ruiz A. Auclair K. Small Molecule Restores Itaconate Sensitivity in Salmonella enterica: A Potential New Approach to Treating Bacterial Infections. Chembiochem. 2016;17:1513–7.

21. Zhang X. Goncalves R. Mosser D.M. The isolation and characterization of murine macrophages. Curr Protoc Immunol. 2008;Chapter 14:Unit-14.1.

22. Vladoianu I.R. Chang H.R. Pechere J.C. Expression of host resistance to Salmonella typhi and Salmonella typhimurium: bacterial survival within macrophages of murine and human origin. Microb Pathog. 1990;8:83–90.

23. Swain A. et al. Comparative evaluation of itaconate and its derivatives reveals divergent inflammasome and type I interferon regulation in macrophages. Nat Metab. 2020.

24. Bocchini V. et al. An immortalized cell line expresses properties of activated microglial cells. J Neurosci Res. 1992;31:616–21.

25. Bambouskova M. et al. Electrophilic properties of itaconate and derivatives regulate the IκBζ– ATF3 inflammatory axis. Nature. 2018;556:501–4.

26. Michelucci A. et al. Immune-responsive gene 1 protein links metabolism to immunity by catalyzing itaconic acid production. Proc Nat Acad Sci USA. 2013;110:7820–5.

27. Duncan D. Auclair K. The coenzyme A biosynthetic pathway: A new tool for prodrug bioactivation. Arch Biochem Biophys. 2019;672:108069.

28. Spry C. Kirk K. Saliba KJ. Coenzyme A biosynthesis: an antimicrobial drug target. FEMS Microbiol Rev. 2008;32:56–106.

29. Fink SL. Cookson BT. Pyroptosis and host cell death responses during Salmonella infection. Cell Microbiol. 2007;9:2562–70.

30. Stapels DAC. et al. Salmonella persisters undermine host immune defenses during antibiotic treatment. Science. 2018;362:1156–60.

31. Barnard L. Mostert KJ. van Otterlo WAL. Strauss E. Developing Pantetheinase-Resistant Pantothenamide Antibacterials: Structural Modification Impacts on PanK Interaction and Mode of Action. ACS Infect Dis. 2018;4:736–43.

32. Howieson VM. et al. Triazole Substitution of a Labile Amide Bond Stabilizes Pantothenamides and Improves Their Antiplasmodial Potency. Antimicrob Agents Chemother. 2016;60:7146–52.

33. Guan J. et al. Structure-Activity Relationships of Antiplasmodial Pantothenamide Analogues Reveal a New Way by Which Triazoles Mimic Amide Bonds. ChemMedChem. 2018;13:2677–83.

34. Jansen PAM. et al. Stable pantothenamide bioisosteres: novel antibiotics for Gram-positive bacteria. J Antibiot. 2019;72:682–92

35. Schalkwijk J. et al. Antimalarial pantothenamide metabolites target acetyl-coenzyme A biosynthesis in Plasmodium falciparum. Sci Transl Med. 2019;11:510.

36. Guan J. et al. A cross-metathesis approach to novel pantothenamide derivatives. Beilstein J Org Chem. 2016;12:963–8.

37. Hoegl A. et al. Stereochemical modification of geminal dialkyl substituents on pantothenamides alters antimicrobial activity. Bioorg Med Chem Lett. 2014;24:3274–7.

38. de Villiers M. et al. Antiplasmodial Mode of Action of Pantothenamides: Pantothenate Kinase Serves as a Metabolic Activator Not as a Target. ACS Infect Dis. 2017;3:527–41.

39. de Villiers M. et al. Structural modification of pantothenamides counteracts degradation by pantetheinase and improves antiplasmodial activity. ACS Med Chem Lett. 2013;4:784–9.

